# Stochastic resonance in the impact of afferent sensory noise on grid-patterned firing and path integration in a continuous attractor network

**DOI:** 10.1101/2024.09.19.613994

**Authors:** Harshith Nagaraj, Rishikesh Narayanan

**Author notes:** **Corresponding Author** Rishikesh Narayanan, Molecular Biophysics Unit, Indian Institute of Science, Bangalore 560 012, India., **e-mail:**. **AUTHOR CONTRIBUTIONS** H. N. and R. N. designed experiments; H. N. performed experiments; H. N. analyzed data; H. N. and R. N. wrote the paper. **COMPETING INTEREST STATEMENT** The authors declare that they have no competing interests.

## Abstract

The continuous attractor network (CAN) model has been effective in explaining grid-patterned firing in the rodent medial entorhinal cortex, with strong lines of experimental evidence and widespread utilities in understanding path integration. A surprising lacuna in CAN analyses is the lack of quantitative studies assessing the impact of sensory noise in the velocity inputs on the emergence of grid-patterned activity and path integration. In addressing this, we employed an established 2D CAN model which received velocity inputs from a virtual animal traversing a 2D arena to generate grid fields. We introduced different levels of Gaussian noise to the afferent velocity inputs to the model and assessed its impact on the network functions in generating grid fields and performing path-integration. We computed a position estimate at each time step using the network activity and quantified position accuracy using the difference between the real and estimated positions. We performed all simulations using several trajectories for the virtual animal, as computed grid scores and position accuracy showed pronounced trajectory-to-trajectory variability even in noise-free cases. We found that the presence of low levels of sensory noise was beneficial to the generation of grid fields, specifically with trajectories where there was no grid-patterned activity in a no-noise scenario. With trajectories where there was grid-patterned activity in the absence of noise, low levels of noise improved position estimation accuracy. In contrast, high levels of sensory noise impaired position estimates as well as grid-patterned activity, although position estimates were more sensitive to sensory noise compared to grid-patterned activity. Together, these analyses demonstrate the manifestation of stochastic resonance in a 2D CAN model, where low levels of sensory noise were beneficial towards the emergence of grid-patterned firing and in tracking position. Next, motivated by the proposed role of border-cells as an error-correction mechanism, we introduced north and east border cells and connected them to grid cells based on co-activity patterns. For different levels of noise, we computed grid scores and position accuracy in the presence *vs*. absence of border cells. Interestingly, while border inputs led to grid field formation in cases where grid fields were not generated, their presence only had a marginal impact on position accuracy. Together, our analyses suggest that biological CANs could evolve to yield optimal performance in the presence of noise in biological sensory systems, serving as a stabilizing factor yielding functional robustness through the manifestation of stochastic resonance.

## Introduction

The theory of path integration is one of the influential theories used to explain the phenomenon of spatial navigation, a behavior crucial to the survival of most organisms. It relies on tracking of self-motion and iterative updating of an internal coordinate to track position. The continuous attractor network (CAN) forms one of the widely used frameworks to implement path integration. Here, a network of neurons embedded with strong recurrent connectivity, coupled with asymmetry in the form of head-direction preferences integrates self-motion inputs to represent different spatial locations in the form of different activity states (Burak & Fiete, 2009; Dong & Fiete, 2024; Fuhs & Touretzky, 2006; Khona & Fiete, 2022; Samsonovich & McNaughton, 1997; Savelli & Knierim, 2019). The model explains the formation of grid cells, which are cells that have spatial firing fields which follow a hexagonal firing pattern (Hafting, Fyhn, Molden, Moser, & Moser, 2005). There are lines of evidence that support the toroidal nature of connectivity employed by most versions of the model (Dong & Fiete, 2024; Gardner, et al., 2022; Khona & Fiete, 2022).

Modeling studies have incorporated empirical observations such as phase precession and theta-dependence in grid cell networks (Bush & Burgess, 2014; Hasselmo & Brandon, 2012; Navratilova, Giocomo, Fellous, Hasselmo, & McNaughton, 2012), intrinsic theta resonance as well as heterogeneities in intrinsic properties and synaptic connectivity (Mittal & Narayanan, 2021), after-spike dynamics (Navratilova, et al., 2012), and stochastic spiking (Burak & Fiete, 2009) to examine the impact grid cell networks and their stability. However, these studies feed deterministic self-motion inputs to the system, thereby precluding analyses of the impact of stochasticity in velocity inputs on network performance. Given the ubiquitous nature of noise in biological systems in general, and sensory systems in particular (Faisal, Selen, & Wolpert, 2008), in this study, we address the impact of introducing sensory noise in the afferent inputs to the continuous attractor network. We asked if the introduction of sensory noise would (a) affect the ability of the network to yield stable grid-fields; and (b) alter the ability of the network to accurately track the position of the animal. We used a well-studied 2D CAN model to address these questions through separate quantitative metrics to assess grid-cell stability and the efficacy of path integration across multiple trajectories.

We found that the ability of the continuous attractor network to yield grid fields was extremely robust to the addition of sensory noise, with stable grid fields formed across a range of afferent noise levels. Grid activity was disrupted only at extremely high levels of noise. We observed pronounced trajectory-to-trajectory variability in the measured grid-patterned activity of individual neurons and associated grid scores. Interestingly, with certain trajectories, the model did not yield grid-patterns in a noise-free scenario but adding low levels of noise helped in the emergence of grid fields. In models which yielded grids in a noise-free situation, we examined the impact of noise on path integration. We tracked the position of the virtual animal using the shift in network activity at each time point and computed deviation in the integrated path for networks receiving different levels of afferent noise. We found that low levels of noise reduced the deviation in integrated path, while higher noise levels were detrimental to the efficacy of path integration. These observations indicate that low levels of sensory noise might be beneficial to a 2D CAN model, a phenomenon termed as stochastic resonance.

While low levels of sensory noise could have a beneficial impact on the network, it is plausible that extrinsic inputs could have similar effects. Several studies have posited that external inputs in the form of border cell inputs can lead to stabilization of grid fields and reduction in path-integration error by recalibrating the system (Cheung, Ball, Milford, Wyeth, & Wiles, 2012; Lisa M Giocomo, 2016; Hardcastle, Ganguli, & Giocomo, 2015; Pollock, Desai, Wei, & Balasubramanian, 2018). Motivated by these, we examined the effect of introducing border cell inputs, either as inhibitory or excitatory inputs to the network, on network performance at various levels of afferent noise. In cases where grid fields were absent in the noise-free network devoid of extrinsic border inputs, the addition of border cell inputs resulted in the generation of grid fields. With reference to path integration, border inputs did not enhance the ability of network activity to predict position at low noise levels but led to modest enhancement in prediction when noise levels were medium range. The introduction of border inputs was not sufficient to alleviate the deleterious impact of very high levels of noise on grid-patterned firing or on location estimates.

Together, our results unveil a stabilizing influence of sensory noise on the formation of grid fields, as well as its beneficial impact on the path integration efficacy of the system. Our results also show that while extrinsic border inputs may serve to stabilize grid fields, they have a modest impact on path integration. Our analyses also reveal trajectory-to-trajectory variability in position estimate evolution, emphasizing the need for multi-trajectory analyses in assessing path integration in CAN models.

## Methods

### Development of virtual trajectory

To reduce computational cost, we modified an algorithm that yields virtual rodent movement (Mittal & Narayanan, 2021) to yield faster trajectories in square arena (2 m × 2 m):

- The starting position of the animal was set to be the center of the arena (*x*_0_ = 1, *y*_0_ = 1).
- For each time step (= 1 ms), the heading direction (*ϕ*_*t*_) as well as the distance run (*d*_*t*_) were picked randomly. Thus, the position of the rat at time *t*, (*x*_*t*_, *y*_*t*_) was given by:

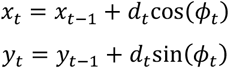

where (*x*_*t*−1_, *y*_*t*−1_) represented the position at the previous time step *t* − 1. If (*x*_*t*_, *y*_*t*_) exceeded the bounds of the arena, they were computed until a random repick of *ϕ*_*t*_ and *d*_*t*_ allowed the coordinates to fall within the arena.
- To make the virtual trajectory more realistic, the range over which the heading direction (*ϕ*_*t*_) and distance run (*d*_*t*_) were picked varied near the boundaries compared to the rest of the arena. To account for the fact that the virtual animal makes fast and sharp turns near the boundaries, the range for *ϕ*_*t*_ was [0, 2π] and *d*_*t*_ range was [0.002, 0.004] m. These ranges were modified to *ϕ*_*t*_ ∈ [–π/36, π/36] and *d*_*t*_ ∈ [0.0029, 0.0031] away from the borders, to achieve smooth runs with relatively uniform speed.

The range of distances per time step (0.002–0.004 m or 0.0029–0.0031 m) and the value of the integration time step (1 ms) implied that the range of velocity was 2–4 or 2.9–3.1 m/s. This implied an average velocity (μ_*v*_) of the virtual animal at 3 m/s across the arena. We generated 50 virtual trajectories, with different virtual trajectories achieved by altering the seed values of the random number generators used for generating *ϕ*_*t*_ and *d*_*t*_. These virtual trajectories offered better control over simulation periods and better efficacy in covering the entire arena in short time spans.

### Structure and dynamics of the continuous attractor network

The continuous attractor model used here was adapted from a previous study (Burak & Fiete, 2009). We used a network of 3600 neurons, which were arranged in a 60 × 60 lattice. The network had periodic boundary conditions, which led to a toroidal arrangement. The neurons in the network were modeled as integrators, with a membrane time constant τ_*m*_ (default value: 10 ms). Each neuron had a preferred direction *θ*_*i*_, implying that they received inputs from specific head-direction cells. *θ*_*i*_ could take one of four different values: 0, π/2, π and 3π/2 which representing the directions east, north, west, and south respectively. The network was divided into local blocks of 2 × 2 cells representing each of these directions (the directional preference of each cell was picked randomly). Thus, one-fourth of the total number of cells in the network preferred a particular direction.

Each neuron in the model received two sets of inputs: an intra-network recurrent inputs from other neurons in the 2D neural sheet and feed-forward inputs that carried information on animal velocity and head-direction preference. The intra-network connectivity followed a shifted Mexican-hat connectivity pattern, which was achieved as a difference of Gaussians, given by (Burak & Fiete, 2009):

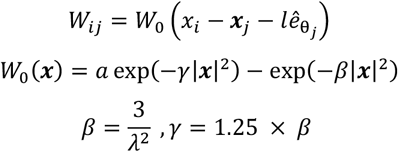

where, *W*_*ij*_ represented the synaptic weight from the neuron located at ***x***_*j*_ to the neuron located at ***x***_*i*_ and 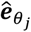 depicted the unit vector pointing along the *θ*_*j*_ direction. The parameter *a* regulates the synaptic weights, and was set to a default value of 1, which would represent an all-inhibitory network. The parameters *β, λ (*default value: 13), and *γ* defined the periodicity of the lattice, while the parameter *l* (default value: 2) defined the shift along 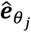, and is necessary for the motion of the activity bump consequent to velocity inputs.

The second set of inputs were feed-forward velocity inputs (based on head direction preference) and were excitatory in nature. These were of the form:

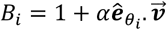

where, *α* (default value: 45) denoted the velocity gain, with 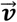 representing the velocity vector derived from the virtual trajectory of the animal.

The dynamics of each of the 3600 rate-based integrator neurons was governed by:

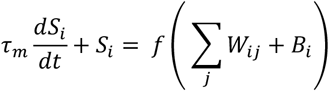

where *f*(*x*) represented the neural transfer function as a simple rectification non-linearity (*f*(*x*) = max(*x, ε*); *ε* = 0), and *S*_*i*_(*t*) denoted the activity of neuron *i* at time *t*. The models were initialized with randomized values of *S*_*i*_ for all neurons with 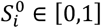. For the initial 100 ms, a constant feed-forward drive was provided to the network by ensuring velocity input was zero, and this resulted in the spontaneous emergence of a stable pattern. The time step (*dt*) was 1 ms and activity patterns were recorded at each time step. For visualization purposes, a spike threshold of 0.1 (a.u.) on *S*_*i*_ was used.

### Introduction of afferent sensory noise

We incorporated sensory noise into the system in the form an additive perturbation 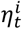 to the velocity vector 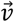 for each cell *i* at each time step *t*. Thus, the feed-forward input to a neuron *i* was now given by:

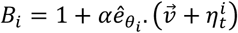

where 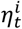 was picked from a Gaussian distribution with zero mean and standard deviation σ _η_. The six levels of noise (L0–L5) were achieved by varying σ _η_ *=* 0, 1×10^−3^, 2×10^−3^, 3×10^−3^, 5×10^−3^, and 1×10^−2^ (L0–L5, in that order).

### Quantitative analysis of grid cell activity

To quantitatively assess the impact of sensory noise on grid-cell activity, we used the following set of standard measurements of grid-cell activity (Fyhn, Molden, Witter, Moser, & Moser, 2004; Hafting, et al., 2005; Mittal & Narayanan, 2021). To quantify the rate maps of grid-cell activity in the network, we first divided the arena into 100 × 100 pixels. *Activity spatial maps* were constructed for each cell by taking the activity of the cell at each time and assigning the activity the cell to the specific (*x, y*) location of the animal in the arena at that time point. *Occupancy spatial maps* were constructed by computing the probability (*p*_*m*_) of finding the animal in the *m*^*th*^ pixel as the fraction of time spent by the animal in the corresponding pixel. *Spatial rate maps* for each cell were computed by normalizing the respective activity spatial map with the occupancy spatial map for that run. These maps were smoothened using a 2D Gaussian filter of standard deviation of 4 pixels. *Average firing rate* (μ) was computed by summing the activity of all the pixels from the spatial rate map and dividing the quantity by the total number of pixels (10,000). *Peak firing rate* (*f*_*m*ax_) denoted the highest activity value observed in the spatial rate map.

To identify grid fields, all local maxima in the smoothened spatial rate map were detected and individual fields were identified as contiguous regions around the local maxima, with a spatial spread of at least 20% activity relative to the local maxima. The *number of grid fields* was computed as the total number of local peaks detected in the spatial rate map of each cell. The *mean size* of each field was estimated by calculating the ratio between total number of pixels covered by all grid fields to the number of grid fields. The *average spacing* between grid fields was the ratio of the sum of distances between the local maxima to the total number of distance values.

*Grid score* was computed by assessing the rotational symmetry of the spatial rate maps. For each neuron, the spatial auto correlation (*SAC*_0_) was computed between the spatial rate map and the map rotated by ϕ° for ϕ ∈ [30, 60, 90, 120, 150]. These values were used to obtain the grid score as follows:

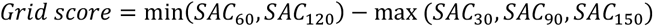

*Spatial information rate (I_S_)* in bits per second was calculated as:

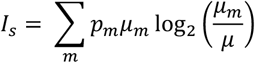

where, *p*_*m*_ defined the probability of finding the animal in the *m*^*th*^ pixel, *μ*_*m*_ denoted the mean firing rate of the cell in the *m*^*th*^ pixel and *μ* was the average firing rate of the grid cell. *Sparsity* was computed as:

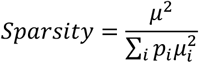

where, *μ*_*i*_ represents the mean firing rate of the cell in the *i*^*th*^ pixel, and *p*_*i*_ represents the probability of finding the animal in the *i*^*th*^ pixel. A lower value of sparsity indicates that the grid-fields are spatially denser.

### Quantitative analysis of path integration

To arrive at a quantitative estimate of path integration, we attempted to arrive at an estimate of the current location of the animal based on network activity (Fig. 1*C–D*). To do this, we first computed the *activity difference map*, defined as the difference between the network activity (activity of each of the 3600 neurons represented as a 60 × 60 matrix) at time (*t* + 1) and time *t*. This is equivalent to computing the differential of the network activity and would yield the ‘shift’ in network activity between time *t* and (*t* + 1). The activity difference map was smoothed with a 2D Gaussian filter with a standard deviation of 1. We estimated the locations of the maxima and minima of the activity difference map, with the maxima denoting cells more active at time (*t* + 1) compared to time *t*, while the minima denoted the cells more active at time *t* compared to time (*t* + 1). Since these local extrema formed clusters (representing the ‘shift’ in a particular activity bump), independent convex hulls were identified around each local maxima and the local minima. The positions of the centroid for each hull were then computed for all local maxima and minima. The differences between the centroids of the adjacent local maximum and the corresponding local minimum were computed for each pair. These differences were averaged across pairs at each time point to give a measure of the network pattern flow velocity for that time point (Fig. 1*C*).

**Figure 1.**
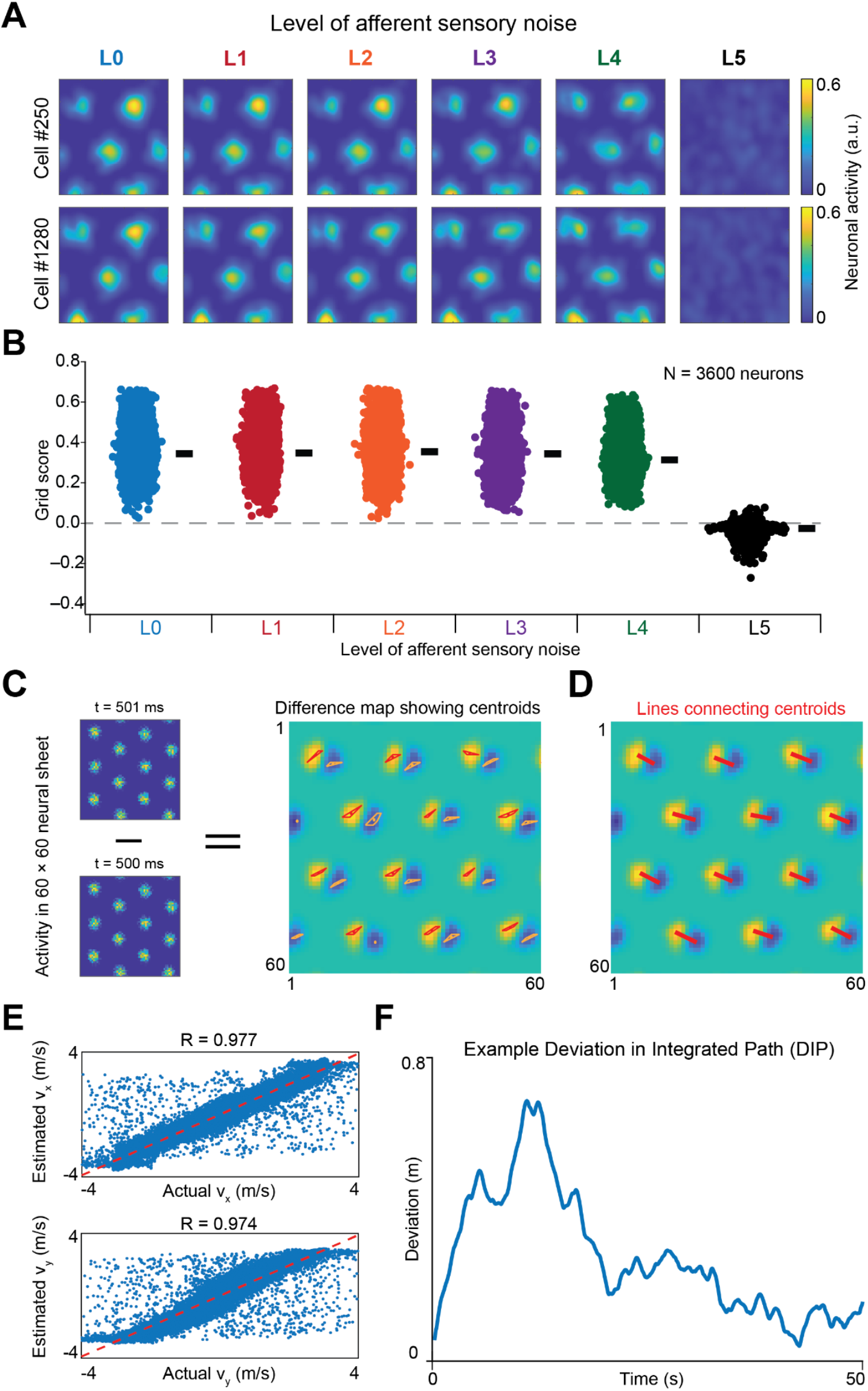
Robustness of grid-patterned firing to the introduction of afferent noise. (*A*) Example spatial rate maps for two representative cells at different levels of noise. The absence of grid-patterned firing may be noted for the map obtained with the highest noise level. (*B*) Bee-swarm plots of the grid score of all 3600 neurons at different levels of noise. The black bar represents the respective median value. Statistical comparisons across different levels of noise are shown in Supplementary Table S1. (*C*) Activity difference map at an example timepoint (500 ms) computed as the difference between the activity map at 501 ms and the activity map at 500 ms. The regions in yellow represent the maxima (thus denoting the cells that were more active at 501 ms), while ones in blue represent the minima (cells that were more active at 500 ms). The convex hulls for the maxima are represented in red, while those for the minima are represented in yellow, with the respective centroids of the computed hulls represented as dots. (*D*) The red lines represent the difference between the centroids of the convex hulls of the adjacent maxima and minima. All red lines are averaged to yield the network pattern velocity, providing the magnitude and direction of shift in the network pattern. The magnitude of the average shift represents the mean hull separation. (*E*) Scatter plots representing the relationship between the *v*_*x*_ (top) and *v*_*y*_ (bottom) components of the estimated *vs*. the actual velocity values computed across all time points. (*F*) Deviation in integrated path, *D*_*IP*_, plotted as a function of time for an example run.

The network pattern flow velocity (computed from the average shift in network activity) would be proportional to the corresponding velocity input to the network 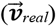 at each time point. The linear fit between the average shift and 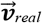 was computed across all timepoints. The average shift was then scaled by the slope of the linear fit to yield the estimate of velocity using the network activity 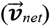. In other words, 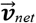 was obtained by scaling the network pattern flow velocity whereas 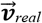 depicted the velocity of the virtual animal in the arena. We computed 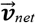 for all time points after 100 ms (as the velocity input was zero for 100 ms for the spontaneous emergence of stable activity patterns).

As the initial position of the animal at 100 ms was known (*x*_100_ = *x*_0_ = 1; *y*_100_ = *y*_0_ = 1), we computed the position estimate of the 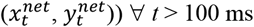 using 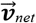 as:

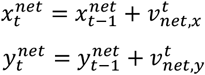

Note that 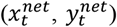 is an *estimate* of the position of the animal computed using network activity. This estimate could be different from (*x*_*t*_, *y*_*t*_), the *actual* position of the animal based on the virtual trajectory. To quantify the difference between the position estimate obtained using network activity 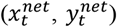 and the actual position (*x*_*t*_, *y*_*t*_), we defined *deviation in integrated path, D*_*IP*_ as follows:

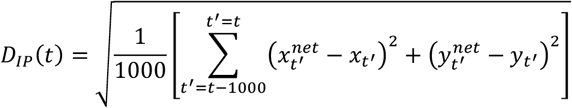

In words, *D*_*IP*_ was computed as the square-root of the sum of the average squared-difference in the *x* and *y* coordinates over a period of 1 s, computed for all time points. The square term ensured sign-independence while the average over 1 s minimized the effect of transient fluctuations. Thus, *D*_*IP*_ provided a quantitative measure of the accuracy with which we can estimate the position of the animal using its network activity.

### Introduction of border cell inputs

A border cell was defined as a cell which manifested high activity exclusively near the border of the arena. We incorporated two different border cell inputs, with high activity either near the north border or along the east border. We chose the two border cell inputs in this fashion, with the hypothesis that the north border cell would correct for errors along the *y* direction whereas the east border cell would correct for errors along the *x* direction. This formulation enabled a scenario where we could examine the impact of border cells with the introduction of a minimal number of additional inputs, apart from providing control over the sign (excitatory *vs*. inhibitory) and magnitude of inputs arriving through these additional border cells to the grid cell network. The spatial firing pattern of the north (*B*_*N*_) and the east (*B*_*E*_) border cells were defined as follows:

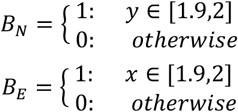

We noted that this formulation for border cell firing to be within 10 cm close to the border is consistent with the firing pattern of border cells observed empirically (Savelli, Yoganarasimha, & Knierim, 2008; Solstad, Boccara, Kropff, Moser, & Moser, 2008) and with previous computational studies (Hardcastle, et al., 2015).

While previous studies have used experience-dependent Hebbian learning to construct the weights of border cell inputs to grid cells (Pollock, et al., 2018), we adopt the approach used by (Hardcastle, et al., 2015) wherein we assume the presence of mature connections between border and grid cells. Specifically, we use network activity obtained from one of the virtual trajectories to construct the connectivity map between border and grid cells. We constructed two different types of connectivity maps between border and grid cells using this process as follows:

- **Excitatory connectivity:** Considering *R*_*d*_ (*d* ∈ {*N, E*}) to denote the region over which the border cell *B*_*d*_ was active, we defined the strength of connection between the border cell *B*_*d*_ and a grid cell *i* (*W*_*id*_) to be proportional to the sum of the activity of the grid cell *i* over the region *R*_*d*_:

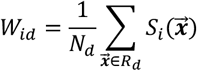

where, 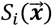 represented the activity of grid cell *i* at position 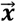 and *N*_*d*_ was a scaling factor (default value: 10,000). The scaling factor was set such that the strength of border-to-grid connectivity was comparable to the strength of the Mexican hat connectivity. This formulation yielded excitatory connectivity, whereby grid cells that were active over the region where the border cell was active received excitatory connections from the border cell. Grid cells that were coactive with the border cells received excitatory inputs from border cells with the strength of excitation directly related to the level of co-activation.
- **Inhibitory connectivity:** For inhibitory connectivity, *W*_*id*_ was defined (with the same notations as above) as:

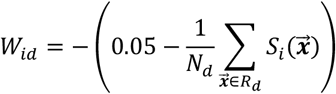

With this formulations, the connections between the border cells and the grid cells are inhibitory in nature. The grid cells that were active over the region *R*_*d*_ received less inhibition compared to grid cells that were not active in the region *R*_*d*_. In other words, all grid cells received inhibition from border cells, but grid cells that were co-active with border cells received reduced inhibition. The magnitude of reduction in inhibition was directly dependent on the level of co-activation.

With the introduction of border cell inputs, the dynamics of the individual cells of the continuous attractor network was modified as:

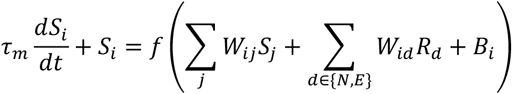

with the additional term involving *R*_*d*_ representing the two border cell inputs.

### Computational details

All simulations were performed in MATLAB 2021a (Mathworks Inc, USA) with a simulation step size of 1 ms. All data analyses and plotting were performed using custom-written software within the IGOR Pro (Wavemetrics, USA) or MATLAB environments. All statistical analyses were performed using the R statistical package (http://www.r-project.org/). To avoid false interpretations and to emphasize heterogeneities in simulation outcomes, the entire range of measurements are reported in figures rather than providing only summary statistics (Marder & Taylor, 2011; Mittal & Narayanan, 2021; Rathour & Narayanan, 2019).

## Results

The goal of this study was to assess the impact of afferent sensory noise and additional extrinsic inputs (in the form of border cells) on CAN performance using different quantitative metrics. We implemented a rate-based CAN model made of a 2D neural sheet (60 × 60 = 3600 neurons) built with integrator neurons (Burak & Fiete, 2009). We introduced afferent sensory noise to the feedforward input and examined its impact on the network.

### Robustness of grid-patterned activity to afferent sensory noise

Existing analyses of grid-cell CAN models do not account for the presence of noise in the afferent inputs to the network, which is ubiquitous across all sensory modalities. Would incorporation of stochasticity in afferent inputs affect the ability of the continuous attractor network to yield stable grid fields? How robust is the network to the addition of noise in afferent sensory inputs? To address these questions, we introduced noise in the afferent inputs to the 2D CAN in the form of an additive Gaussian of zero mean and variable standard deviation (σ_η_). The standard deviation of the Gaussian was varied to yield 5 different levels of noise (L1–L5) with L0 denoting the deterministic case where no noise was introduced. Neural activity was recorded as a virtual animal traversed the square arena. The activity map of the 3600 neurons in the network as a function of 2D space within the arena was computed across different levels of noise.

Example patterns of activity across the arena show that grid-patterned activity was remarkably robust to the addition of noise (Fig. 1*A*). To quantify this across all neurons in the network, we computed grid score for each neuron in network. The grid scores (Fig. 1*B*) and other grid-field measurements (Supplementary Fig. S1*A–G*) confirmed the remarkable robustness of grid-patterned firing to the addition of sensory noise. Specifically, we found stable grid-fields across most noise levels, with the manifestation of instability limited to the highest level of noise (Fig. 1; Supplementary Fig. S1). The design of virtual trajectory was such that the mean velocity of the animal during the run (μ_*v*_) was 3 × 10^−2^ m per time step (1 ms). The remarkable nature of the robustness of grid-patterned firing to noise is evident, as stable grid patterns were observed even when the ratio σ_η_/μ_*v*_ exceeded 1. The disruption in grid-patterned firing occurred at L5 level of noise where σ_η_/μ_*v*_ was greater than 5.

### Quantifying the accuracy with which the 2D CAN tracks position

How does the estimate of animal position obtained from the network activity compare with the actual position as a function of time? To characterize the ability of network activity to track animal position and to assess the impact of noise on the accuracy of the position estimate, we defined a metric termed as the *deviation in integrated path, D*_*IP*_(*t*). *D*_*IP*_ (*t*) was computed over a period of 1 s for all time points *t*, as the average squared difference between the position estimate (obtained from the network pattern flow velocity) and the real position of the virtual animal in the arena. *D*_*IP*_ (*t*) represented the accuracy with which we can estimate the animal’s position using the network activity and track this as the virtual animal traverses the arena. Quantitatively, a lower value of *D*_*IP*_ indicated enhanced accuracy in the position estimate.

In computing *D*_*IP*_, we estimated network pattern flow velocity from the difference maps of network activity obtained at two consecutive time points (Fig. 1*C*). We identified the centroids of the adjacent positive and negative lobes associated with each grid field in the difference map by computing the centroids of independent convex hulls for the two lobes (Fig. 1*D*). The lines connecting the two centroids (associated with positive and negative lobes of a given grid field in the difference map), computed across all grid fields, provided an estimate of network pattern flow velocity at each time point. We scaled the network pattern flow velocity to arrive at a velocity estimate of the animal 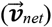 and compared it with the actual velocity of the animal 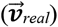. We found 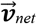 and 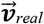 to be highly correlated (Fig. 1*E*). In the network that lacked sensory noise (L0 level), with a given virtual trajectory, we found *D*_*IP*_ to initially increase as a function of time but to eventually settle down at a low level of deviation (Fig. 1*F*).

### Sensory noise-driven stochastic resonance in the generation of grid-patterned activity

How sensitive is the propensity of the CAN to yield grid-fields to the specific trajectory used? Does using a different trajectory (in the same arena) affect the nature of grid fields obtained? To answer this question, we generated 60 different trajectories and assessed grid-patterned activity in the network, each trajectory assessed with 5 different levels of noise. In the noise-free scenario, the network yielded stable grid fields with most trajectories (50/60=83% of trajectories tested). Interestingly, even in such a noise-free scenario, we observed that the network did not generate grid-fields in certain trajectories (Fig. 2*A*, Supplementary Fig S4). Strikingly, in these cases, we found that adding a low level of noise in the afferent velocity inputs to the network resulted in the manifestation of grid-fields, with each of L1–L4 levels noise levels leading to grid field formation (Fig 2*A–C*, Supplementary Fig S4). We quantified the “gridness” of the rate maps by computing the grid score (Methods), which showed a significant increase with the addition of noise (Fig 2*B–C*). The highest level of noise L5, however, led to a complete loss of grid patterned activity (Fig 1*A*). These results demonstrate that adding low levels of sensory noise could be beneficial to the CAN in generating grid-fields, a phenomenon referred to as stochastic resonance, defined as the ability of an optimal level of noise to improve system performance, with performance dropping on either side of the optimal noise level (McDonnell & Abbott, 2009).

**Figure 2.**
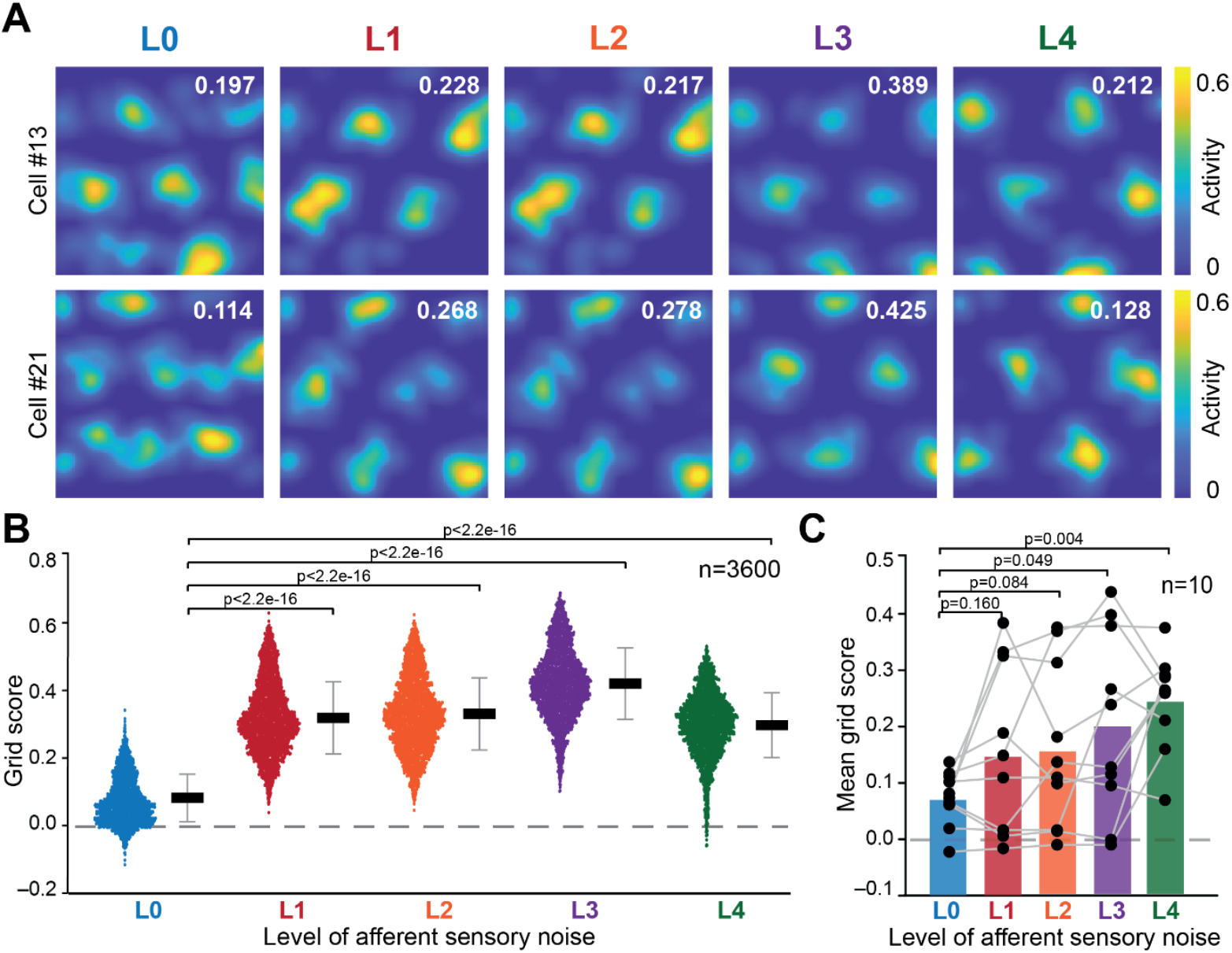
Addition of sensory noise noise is beneficial to the formation of grids. (*A*) Rate maps of two cells for an example trajectory showing the emergence of grid-fields on the addition of sensory noise for an example trajectory. The values in white refer to the grid score of that particular cell. (*B*) Grid scores for each cell across different noise levels for the example trajectory from above. (*C*) Mean grid score across all cells as a function of noise level, across different trajectories (*n* = 10). The black dots and gray lines represent results from individual trajectories.

### Sensory noise-driven stochastic resonance in the accuracy of position estimated from network activity

When the network generates stable grid-fields, what does the addition of sensory noise do to the system’s ability to estimate position? Given the ubiquitous presence of noise in biological systems, is the 2D CAN able to track position accurately in the presence of sensory noise? To answer these questions, we computed *D*_*IP*_ for the 50 trajectories that yielded grid-fields in a noise-free scenario and examined the effects of adding noise. As the identification of convex hulls is an essential requirement for estimating position from network activity, we were unable to compute *D*_*IP*_ for networks that did not show stable grid-patterned activity.

For a single trajectory, we observed that the addition of noise paradoxically resulted in an increase in the accuracy of the position estimate of the animal obtained using the network velocity. Specifically, compared to L0 noise level, we noticed a reduction in *D*_*IP*_ across most timepoints for the L1 and L2 levels of noise, whereas L3 and L4 levels of noise resulted in an increase in *D*_*IP*_ across most timepoints (Fig. 3*A*). We could not quantify the deviation for L5 levels of noise due to our inability to construct the convex hull (Supplementary Fig. 1*H*) owing to the absence of well-defined grid fields (Fig. 1).

**Figure 3.**
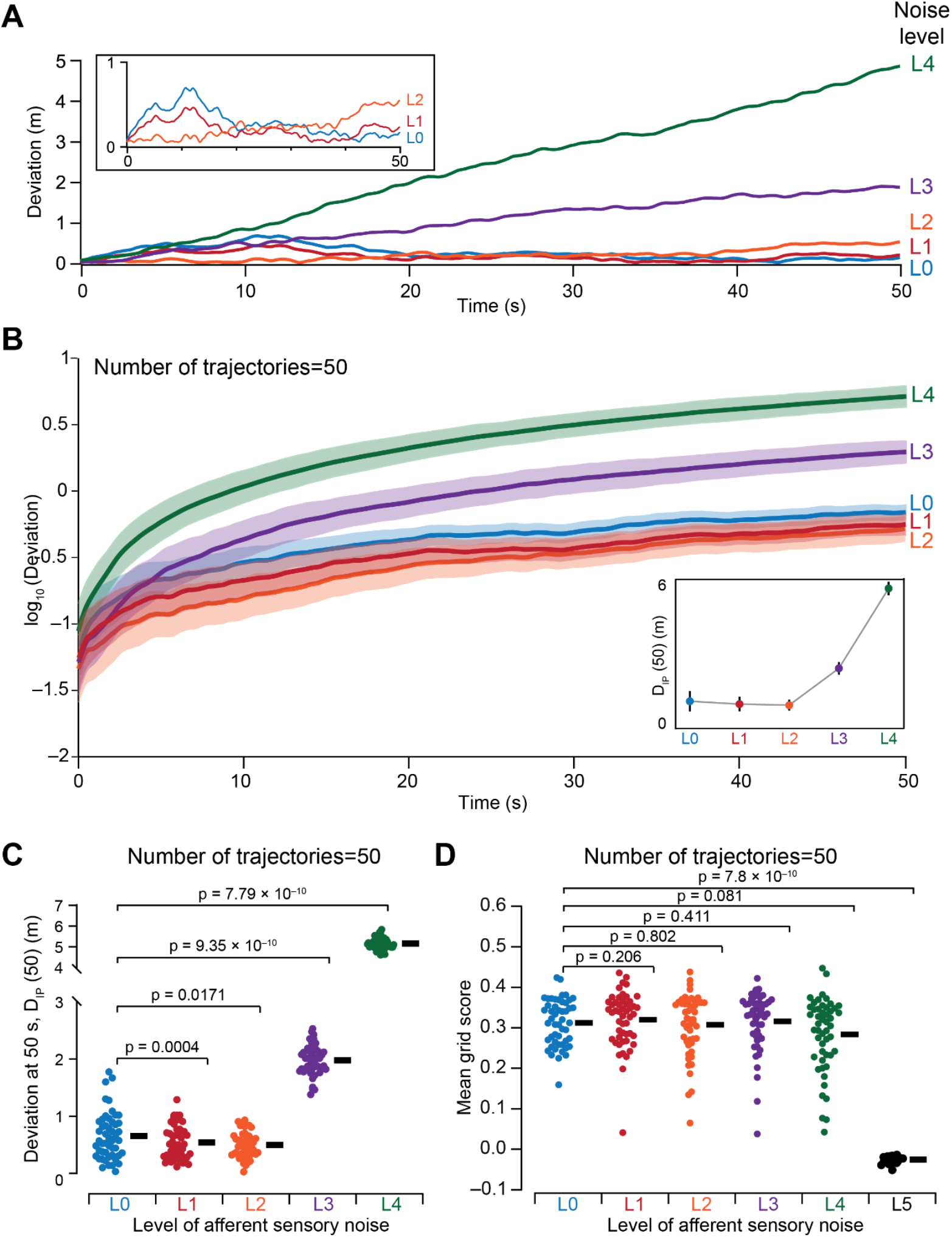
Low levels of afferent noise enhanced position estimates. (*A*) Example traces of deviation in integrated path, *D*_*IP*_, computed for a single trajectory at different levels of noise (L0–L4). (*Inset*) Zoomed version of the traces for L0–L2 levels of noise. (*B*) Mean-SEM of *D*_*IP*_, computed across 50 different virtual trajectories, plotted on log scale for different levels of noise (L0–L4). The solid line represents the mean across the 50 trajectories while the shaded region represents the standard error. (*Inset*) *D*_*IP*_ value at the 50 s timepoint plotted as a function of noise level. Circles represent the mean across 50 trajectories and the bars represent the standard deviation. (*C*) Beeswarm plots representing *D*_*IP*_ value at the 50 s timepoint for 50 different trajectories at different levels of noise. Each circle represents *D*_*IP*_ computed from a different trajectory. The black bars represent the mean across the 50 trajectories. *p-*values show the outcomes of comparison of no-noise measurements with measurements obtained with different levels of noise and were computed using Wilcoxon rank sum test. The Kruskal Wallis test spanning groups across all noise levels yielded a *p* value of 2.2 × 10^−16^. *(D)* Bee-swarm plots representing the mean grid score across the 3600 neurons in the network for each of the 50 different trajectories at different levels of noise. Each circle represents the mean grid score computed from a different trajectory. The black bars represent the median across the 50 trajectories. *p-*values show the outcomes of comparison of no-noise measurements with measurements obtained with different levels of noise and were computed using Wilcoxon rank sum test. The Kruskal Wallis test spanning groups across all noise levels yielded a *p* value of 2.2 × 10^−16^.

As the specific values of *D*_*IP*_ could be reflective of the specific virtual trajectory employed, we repeated our analyses with 50 different virtual trajectories and computed the deviation (Fig. 3*B–C*) as well as grid score (Fig. 3*D*) for each of the 50 runs. Strikingly, we found that the mean deviation obtained for noise levels L1 and L2 was consistently lower than the mean deviation for the noise-free level L0 (Fig. 3*B–C*), while the grid scores remained similar (Fig 3D). However, for L3 and L4, the deviation was higher compared to the other three levels of noise. These results demonstrated that position estimates improved with an optimal level of sensory noise and performance drops on either side of this optimal level of noise (Fig. 3*C*), together unveiling the manifestation of stochastic resonance in the accuracy of position estimation from a 2D CAN.

An important observation about the need for performing simulations with multiple trajectories emerged from trajectory-to-trajectory variability that we observed in the quantitative metrics across different trajectories for the same network. Specifically, pronounced variability in *D*_*IP*_ (Fig. 3*C*) and average grid scores (Fig. 3*D*) across different trajectories strongly emphasized the need to validate network performance with multiple trajectories, even in noise-free conditions (*e*.*g*., L0 variability in Figs. 3*C–D*). If analyses were performed with a single trajectory, the biases associated with that single trajectory drive the conclusions. For instance, stochastic resonance observed in the deviation measurement (Fig. 3*B*) would not have been evident if analyses relied on a single trajectory (Fig. 3*A*). It required multiple runs to eliminate trajectory-driven biases to accurately assess the impact of sensory noise on this measurement.

Together, these results (Figs. 1–3) demonstrate the beneficial impact of afferent sensory noise on position estimation accuracy and grid-patterned activity generation in a continuous attractor network. Low levels of sensory noise helped in the generation of grid-fields and improved position estimation accuracy. In contrast, high levels of sensory noise impaired position estimates as well as grid-patterned activity, although position estimates were more sensitive to sensory noise compared to grid-patterned activity (Figs. 1–3).

### Addition of border inputs did not affect the path integration accuracy of the system

Border cell inputs are known to serve as an error-correcting mechanism for grid cells (Hardcastle, et al., 2015) and serve to minimize the drift in grid patterns over time (Pollock, et al., 2018). Motivated by this, we asked if the addition of border inputs could increase the accuracy of the position estimate obtained using the network activity in the presence of afferent sensory noise. We introduced border inputs to grid cells using two different types of connectivity: (i) an inhibitory pattern, wherein grid cells coactive with border cells received lower levels of inhibition; (ii) and an excitatory pattern, wherein grid cells coactive with border cells received higher levels of excitation. The connectivity pattern was constructed using grid cell co-activity during a representative run and used for later runs to introduce border cells. Two border cells were considered to provide inputs to grid cells: one that fired along the north border and another along the east border, to potentially serve as corrective influences along orthogonal directions (Fig. 4*A*).

**Figure 4.**
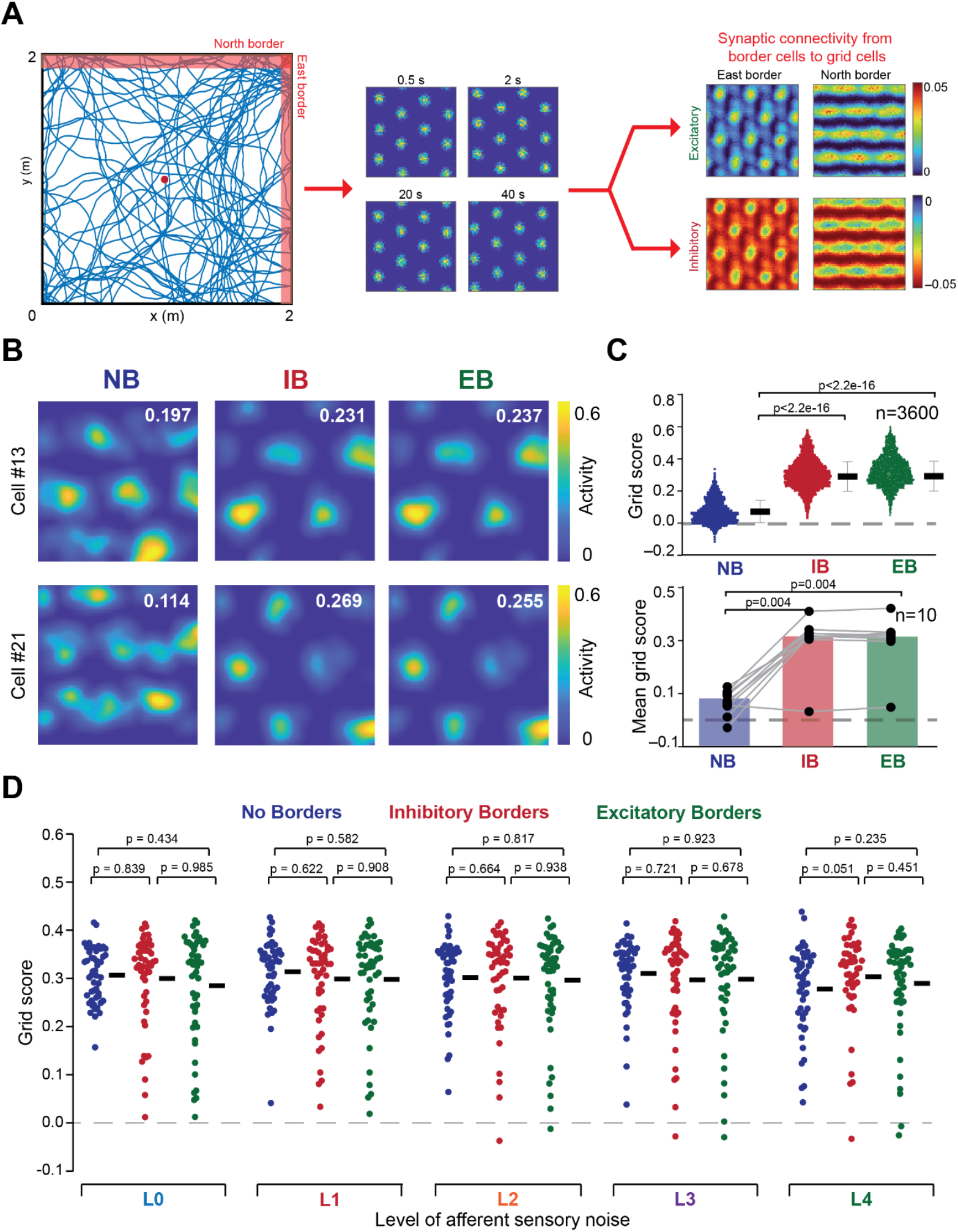
Introduction of border cells, either through excitatory or inhibitory connectivity is beneficial to the generation of grid fields. (*A*) Border-to-grid connectivity maps were constructed using results on co-active grid cells from a representative trajectory. *Left*, 2 m × 2 m virtual arena showing the trajectory used to construct the border-to-grid connectivity maps. Our model consisted of two border cells: one that fired along the north border and one that fired along the east border (red shades in the arena). *Middle*, activity map of the grid neural sheet (60×60) at different time points to figure out grid cells that were co-active when border cells were active. *Right*, Connectivity was either excitatory or inhibitory in nature and was defined by co-activity patterns. (*B*) Rate maps of two grid cells from an example trajectory showing the generation of grid fields on adding border inputs. The values in white refer to the grid score of the individual cells. (*C*) (Top) Grid scores of the 3600 cells in the network for the cases when border inputs were present/absent for the example trajectory in (*B*), and (bottom) Mean gid score across all cells for the different border cases. The black dots and gray lines represent individual trajectories. *p-*values are provided for the Wilcoxon rank-sum test. (*D*) Distribution of mean grid scores (across the 3600 neurons in the network) for the 50 different trajectories, each computed for five different levels of noise (L0–L4). No border results were the same as before. Inhibitory border and excitatory border represent inhibitory and excitatory border-to-grid connectivity patterns, respectively. *p* values are provided for the Wilcoxon rank-sum test. For each level of noise, the *p* values for Kruskal-Wallis test spanning all three border-types were: L0: 0.25; L1: 0.86; L2: 0.73; L3: 0.97; L4: 0.14: L5: 0.50.

For a single trajectory run of simulations, we computed grid-cell metrics for each of the 3600 cells in the absence of noise and found that the addition of border inputs (through excitatory or inhibitory connectivity) had a modest impact on these metrics (Supplementary Fig. S2). To avoid biases associated with the use of a single trajectory, we performed network simulations with border cells over the 60 different trajectories used earlier and with different levels of noise. In cases of trajectories (*n*=10) where there were no grid fields formed in the L0 level, we found that adding border cell inputs led to the formation of grid fields (Fig 4*B–C*, Supplementary Fig S4), with a significant increase in grid scores (Fig 4*C*). This is consistent with existing literature supporting the role of border cell inputs as stabilizing mechanisms to grid fields. However, in the 50 networks that did show grid-patterned firing in the L0 level, we observed no significant increase in grid scores with the addition of border inputs across L0–L4 noise levels (Fig. 4*D*).

We next turned to accuracy of location estimation in the presence of border cells. We had earlier (Fig. 3*C*) established that the presence of low levels of noise enhanced location estimates manifesting as a reduction in the deviation metric *D*_*IP*_ for L1–L2 levels of noise, but increased values of *D*_*IP*_ for L3–L4 levels (all compared to noise-free L0). Under noise-free conditions, the estimation accuracy was statistically comparable irrespective of whether border inputs were present or not (Fig. 5*A*). Strikingly, with low levels of noise (L1–L2), *D*_*IP*_ measured across 50 trajectories was marginally higher when border inputs were present. In contrast, with increase in noise level (L3–L4), the presence of border inputs slightly enhanced the estimation accuracy (significantly so for L4, inhibitory connectivity). However, the *D*_*IP*_ values with L3– L4 noise levels were much higher than in the noise-free scenario, thus demonstrating that adding border inputs doesn’t compensate for the effects of noise at high levels.

**Figure 5.**
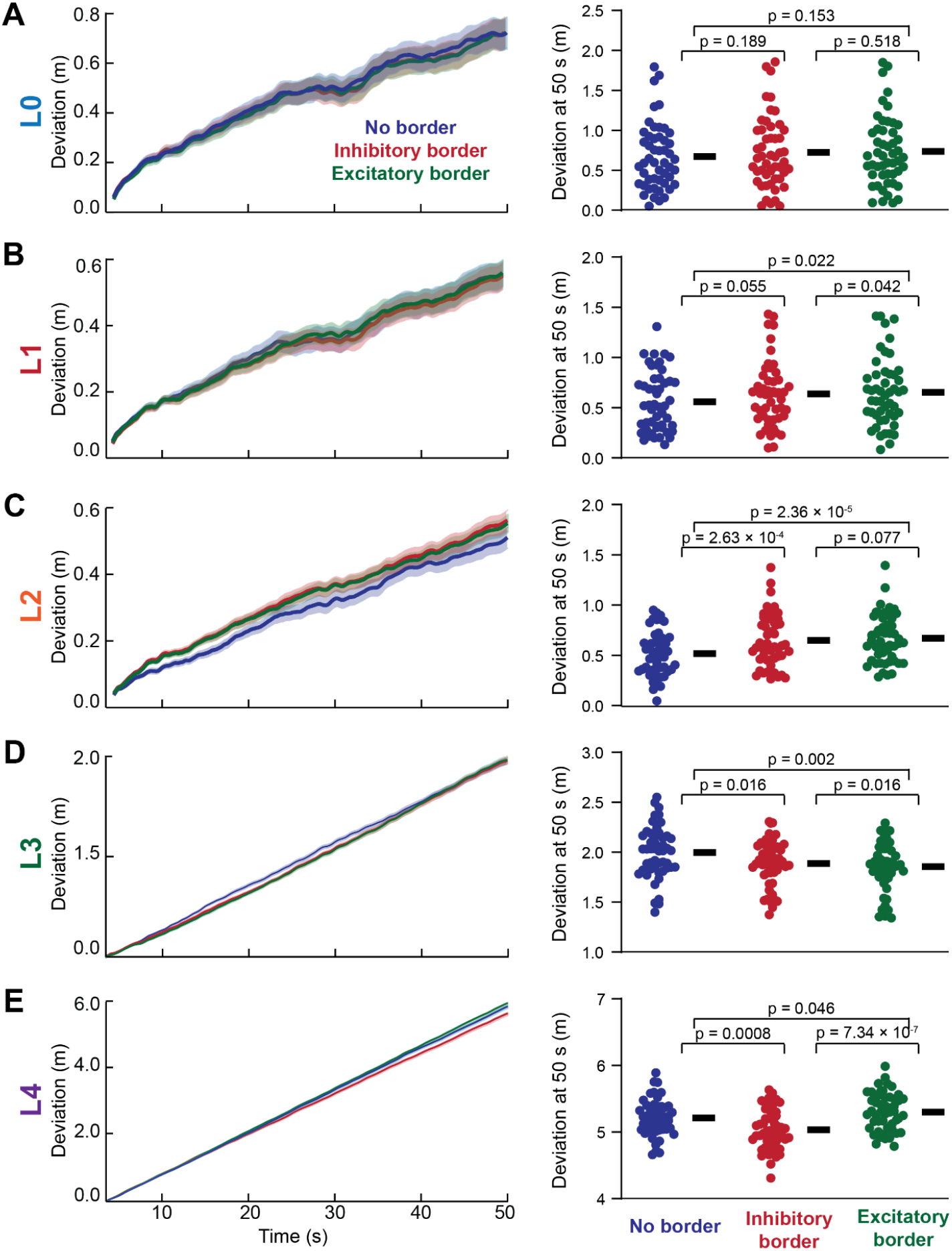
Introduction of border cells, either through excitatory or inhibitory connectivity, yielded modest changes to accuracy of position estimates. (*A–E*) *Left*, Deviation computed across 50 different trajectories as a function of time, for the 3 different types of border inputs (no border, excitatory border, inhibitory border) introduced, for different levels of afferent sensory noise. The solid line represents the mean across the 50 trajectories with the shaded region representing the standard error. *Right*, Bee-swarm plots of the deviation at the 50 s time point for the 50 different trajectories, for 3 types of border input to the network and different levels of sensory noise (L0–L4 spanning panels A–E, respectively). Colors represent the type of border input, with black bars indicating the median. *p* values are provided for the Wilcoxon rank-sum test. For each level of noise, the *p* values for Kruskal-Wallis test spanning all three border-types were: L0: 0.86; L1: 0.60; L2: 3.01 × 10^−2^; L3: 0.20; L4: 2.01 × 10^−5^.

Together, our results show that while the introduction of border inputs led to stabilization of grid-fields in cases where there are no grid-fields formed in the noise-free scenario (Fig 4*B–C*), the addition of border inputs introduced modest differences in position measurements across different levels of noise (Fig. 5). Together with our earlier results (Fig 2*C*, 3*C*), it indicates that sensory noise could play a greater role in stabilizing grid-fields and enhancing the accuracy of position tracking compared to external inputs. They also demonstrate that the introduction of border inputs was not sufficient to alleviate the deleterious impact of high levels of noise on location estimates.

### Progressive reduction in mean hull separation with increase in sensory noise variance contributes to dependence of position estimate on sensory noise

What was the basis for the enhanced position estimates obtained with the low levels of noise (Fig. 3)? To address this question, we revisited the process by which we compute the network pattern flow velocity and how neural activity is related to all inputs (including noisy velocity inputs). The central step in the estimation of network pattern flow velocity is the vector connecting the centroids of the convex hulls constructed for the maxima and minima of the activity difference map (Fig. 1*C, D*). Therefore, we first assessed the impact of afferent noise on this distance (the magnitude of the vector connecting the two convex hulls) and referred to this as hull separation. We computed the distribution of mean hull separation (across all grid fields) at each time point for the same trajectory with different levels of sensory noise. We found that increases in noise levels resulted in a reduction in mean hull separation, manifesting as a progressive leftward shift in the cumulative distribution of mean hull separation with increasing noise level (Fig. 6*A*).

**Figure 6.**
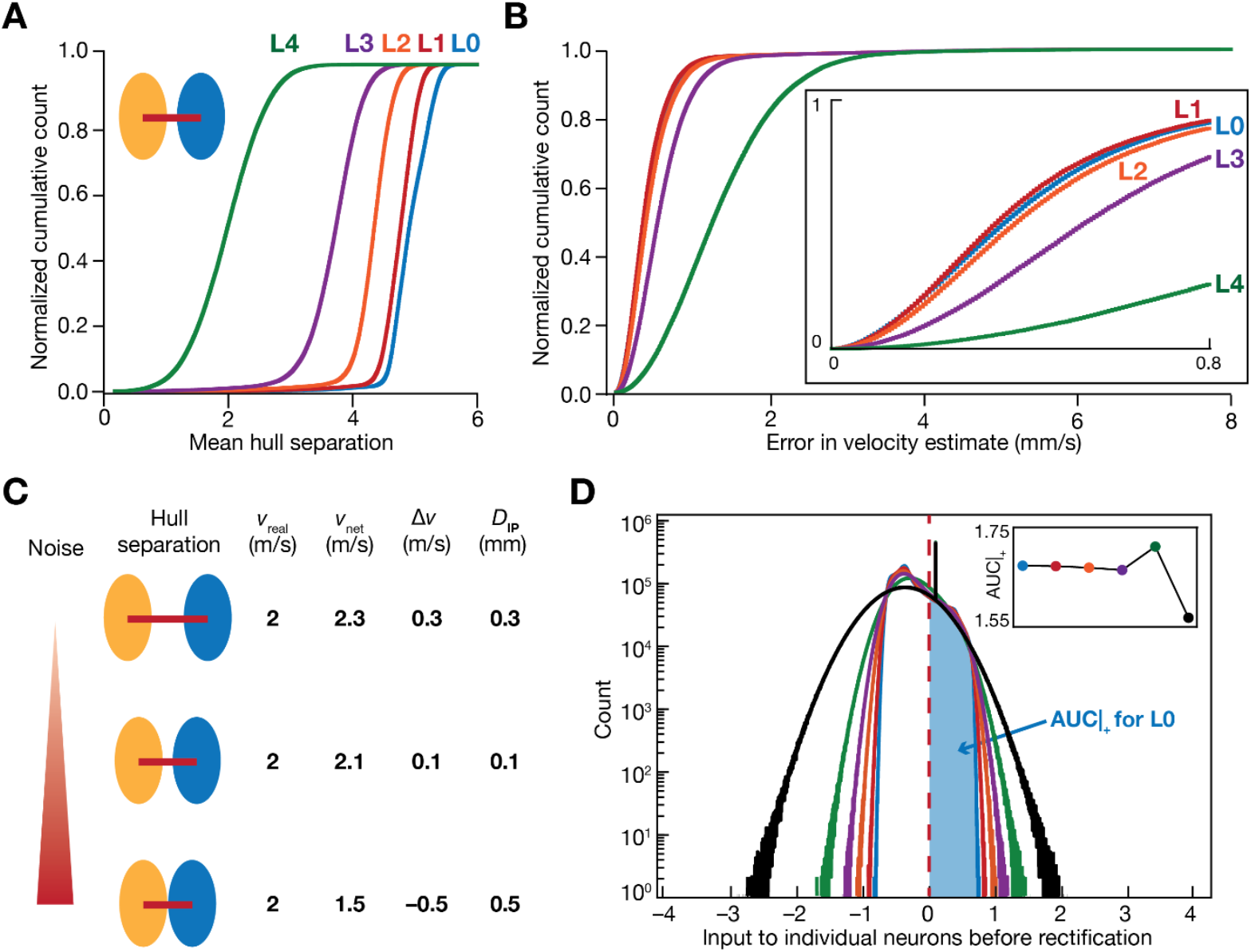
Progressive reduction in mean hull separation with increase in sensory noise variance contributes to dependence of position estimate on sensory noise. (*A*) The yellow and the blue ellipses represent the positive and the negative lobes, respectively, used for computing the network pattern flow velocity (illustrated in Fig. 1*C– D*). The magnitude of the red line connecting the centroids of the two lobes is referred to as hull separation. The mean hull separation for each time point is computed by averaging all values of hull separation across every activity bump. The plots represent the distributions of the mean hull separations for different noise levels (L0– L4). (*B*) Distributions of the error in velocity estimate (difference between the velocity estimated using network activity and the actual velocity) for different noise levels. *Inset*, A zoomed version of the distribution spanning lower values of error. (*C*) Model for how increase in noise reduces hull separation, which in turn could contribute to stochastic resonance in ***D***_***IP***_ observed in Fig. 3*D*. (*D*) Distribution of pre-rectification inputs to the 3600 neurons of the network for a representative trajectory, for different levels of noise. The red line represents the default threshold (***ε***) in the neural transfer function for the input to manifest as neural activity. The shaded blue region represents the area under the curve (AUC) for the positive part of the distribution for L0 noise level. *Inset*, Area under the curve for the positive part of the distributions for L0–L4 noise levels.

Animal position is estimated from the estimated animal velocity from network activity 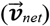, which depends on the hull separation vector. A reduction in hull separation for the same values of real velocity 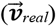 values implies that there would be differences in the animal velocity estimated from network activity. To quantify this, we computed error in velocity estimate (Δ*v*) as the magnitude of the difference between 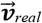 and 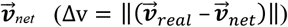 at each time point for the same trajectory with different levels of sensory noise. Owing to the reduction in the hull separation, we observed that the error in velocity estimate was higher in the case of L3 and L4 levels of noise compared to the noise-free scenario (Fig. 6*B*). Notably, we observed that the error in velocity estimates for the noise-free L0 level was slightly higher than the L1 level of noise pointing to hull separation as one potential source for the stochastic resonance observed in *D*_*IP*_ (Fig. 3).

Thus, based on these observations, we propose a model which reconciles the reduction in mean hull separation and the stochastic resonance in *D*_*IP*_ with addition of sensory noise (Fig. 6*C*). In the absence of sensory noise, the mean hull separation is large, which results in an overestimation of the animal’s velocity. This overestimate manifests as a relatively high value of Δ*v* (Fig. 6*B*), resulting in high deviation in zero noise conditions. With increasing noise levels, the mean hull separation progressively reduces (Fig. 6*A*). At an optimal level of low levels of noise, the estimated velocity approaches the real value thus ceasing to be an overestimate, leading to a reduction in *D*_*IP*_ (Fig. 3). Upon further increase in the noise level, the mean hull separation is reduced further (Fig. 6*B*), yielding an underestimation of the real velocity. This manifests as increased error in velocity estimate (Fig. 6*C*) and a higher value of *D*_*IP*_ (Fig. 3).

### The distribution of inputs to the rectifying nonlinearity in individual neurons is a key factor contributing to network estimates of position

What factors contributed to the reduction in hull separation that we had observed with the increase in noise (Fig. 6*A*)? We noted that the rectifying nonlinearity in individual neurons contributes to the conversion of total inputs to neural activity at every time point. We reasoned that increase in noise would enhance the range of the overall inputs arriving onto individual neurons, thus resulting in higher activity in the presence of noise. This higher activity could result in expansion of the lobes at each time point, together yielding a reduction in hull separation. To test this, we recorded inputs onto each neuron in the network across all time points for the same trajectory with different levels of noise. We plotted the distribution of these inputs and found that the input range indeed increased with increasing noise (Fig. 6*D*). These observations indicate that the noise-induced enhancement in range of inputs in conjunction with the rectification resulted in an ability to detect inputs that were sub-zero under no-noise conditions. This enhancement resulted in a reduction in hull separation with increased noise. These observations suggest that the enhancement observed at lower noise levels is not due to an increase in the extent of input range but could be because of noise-induced detection of sub-threshold inputs without alteration to the threshold.

We assessed hull separation in our simulations with border inputs (Figs. 4–5). Consistent with the modest changes in position estimates with the introduction of border inputs, we also found that each of hull separation, velocity estimation error, and pre-rectification input distribution showed modest changes with introduction of border inputs (Supplementary Fig. S3). Overall, the impact of border inputs on position estimation was modest across noise levels and across both kinds of connectivity.

## Discussion

The principal goal of this study was to assess the impact of introducing sensory noise in the afferent inputs to the CAN model on grid-fields and the ability of the network to track position. Our analyses demonstrate the robustness of the grid-firing fields generated by the model to sensory noise, with disruption in patterns observed at extreme levels of noise. We report the manifestation of stochastic resonance in the grid-field emergence and accuracy of position estimation from network activity in the presence of sensory noise. Specifically, we found that low levels of noise were beneficial in grid-field emergence and in estimating the animal position with greater accuracy, while high levels of noise were detrimental. Finally, we found that the introduction of border cell inputs to the grid cell network had modest impact on position estimation while being beneficial to grid-field emergence.

While probing the mechanism behind this stochastic resonance associated with sensory noise, we found that low levels of noise enhanced detection of sub-threshold sensory signals, which in turn enhanced the ability of the resultant activity patterns to enhance accuracy of estimated location. On the other hand, with high levels of noise, the dominance of noise over the velocity signal degraded efficient path integration. Together, this yields an optimal level of noise that leads to minimal error in path integration. However, this enhancement is not observed when border cell inputs were added, which is consistent with the modest impact on path integration.

### Robustness of the grid patterns to sensory noise

Noise is ubiquitous in biological systems across all scales. Sources of noise in biological systems include stochasticity in ion-channel opening, synaptic release, binding of molecules involved in signaling cascades, which can propagate across scales to introduce widespread variability in responses (Faisal, et al., 2008; Ribrault, Sekimoto, & Triller, 2011). Thus, the fundamental requirement for a biological system is to maintain its functionality in the presence of such stochasticity. With reference to CAN models, although the impact of noise in neural spiking has been assessed (Burak & Fiete, 2009), the impact of sensory noise at the input end of the neurons has not been assessed. Our observations on the stability of grid-patterned firing patterns over a wide range of noise levels demonstrate the extent of robustness exhibited by the CAN model. Such robustness in pattern generation is expected from a network exhibiting attractor dynamics, wherein the specific connectivity patterns and stable activity propagation across the neural sheet counteracts the perturbations induced by noise (Cannon, Robinson, & Shamma, 1983; Khona & Fiete, 2022). The presence of multiple ‘activity bumps’ in the neural sheet also has been shown to contribute to robustness to noise (R. Wang & Kang, 2022). Our observations on robustness of grid-patterned firing to a wide range of afferent noise are therefore consistent with prior observations and is a reason for the attractiveness of the framework for patterned activity generation.

### Stochastic resonance in grid-field emergence and in accurate position estimation

Stochastic resonance has been observed in neural systems across scales, ranging from molecules to single neurons to perceptual levels (Collins, Chow, & Imhoff, 1995a, 1995b; Douglass, Wilkens, Pantazelou, & Moss, 1993; Gluckman, et al., 1996; Mittal & Narayanan, 2022; Moss, 1997; Russell, Wilkens, & Moss, 1999; Simonotto, et al., 1997). Our analyses show that the addition of low levels of noise is beneficial to the emergence of grid fields, specifically in cases where the network is unable to generate grid fields in the absence of noise. In cases where grid fields are generated even in a noise-free scenario, we demonstrate the beneficial effect of low levels of noise in estimating the position accurately using network activity, with higher levels of noise resulting in reduced accuracy. The manifestation of stochastic resonance implies that a biological CAN could evolve in the presence of sensory noise, such that grid-patterned firing and optimal path integration are achieved at the specific level of noise that is prevalent in the system. Although we were able to partially explain the reason behind this stochastic resonance using the rectification nonlinearity in the neural activation function, further analyses are required to fully elucidate the underlying mechanisms.

### Modest impact of introducing border cell inputs on path integration

Path integration within the CAN framework relies on the iterative update of an internal coordinate by tracking self-motion cues. Drift in position estimates have been shown to accumulate iteratively under such a framework. While studies have relied on border-cell inputs to act as ‘corrective influences’ under such a scenario (Lisa M Giocomo, 2016; Hardcastle, et al., 2015; Pollock, et al., 2018), our results on the effect of borders on the accuracy of the position estimate under different noise conditions yield important insights. Firstly, border cell inputs were conducive to the generation of grid fields in cases when the network is unable to generate them independently, which is broadly consistent with previous results (Lisa M Giocomo, 2016; Hardcastle, et al., 2015; Pollock, et al., 2018). We also found that border inputs did not seem to significantly improve the accuracy of path integration when compared to low levels of noise, nor were they able to counteract the disruptive effect of noise at high levels. Thus, it implicates a dichotomy in the effect of border inputs: one where they stabilize grid-fields while having marginal impact on the position tracking accuracy.

### Need to account for trajectory-to-trajectory variability

Irrespective of whether the network received border inputs or noise, we observed pronounced trajectory-to-trajectory variability across all measurements. It was therefore essential to compute each measurement across multiple trajectories to avoid biases that are introduced using a single trajectory. With specific reference to path integration and temporally continuous position estimates, the exact nature of each trajectory would result in different errors as a function of time. To study the impact of another variable, such as sensory noise variance or border-cell inputs, on path integration performance, it becomes essential that the dependence of error evolution on individual trajectories is eliminated. Without such elimination, the outcomes are biased by the specific dynamics of that single trajectory thereby providing incorrect inferences. Our analyses demonstrate that the impact of sensory noise (Fig. 3) or border inputs (Figs. 4–5) on network performance showed wide trajectory-to-trajectory variability and inferences about the new variable were decipherable only across 50 different trajectories. Future studies should study the source of trajectory-to-trajectory variability and how they affect the impact of specific variables on overall network performance using different metrics that probe the varied functions.

### Future directions and model limitations

Our study employs a rate-based model of a continuous attractor network, which is endowed with noisy sensory inputs as well as border inputs to generate grid-patterned activity as well as to estimate position using network activity. Rate-based models are limited in their ability to account for temporal characteristics of spike timings as well as the relationship to the phase of extracellular oscillations (such as the theta oscillation), which are necessary for the spatial selectivity of grid cell activity (Brandon, et al., 2011). Future studies expand our analyses using spiking or biophysical neural models that are endowed with other forms of noise including synaptic and channel noise.

Our model employs a homogeneous population of neurons, whereby each neuron possesses similar intrinsic properties and is endowed with similar connectivity patterns. Biological systems, however, are endowed with significant heterogeneity in their properties. These heterogeneities have been shown to beneficial or deleterious effects of network function, depending on several factors including the circuit under consideration, the intrinsic properties of neurons, the connectivity pattern, the task implemented, the form and degree of the heterogeneity, and interactions among different forms of heterogeneities (Brunel & Hakim, 1999; Dahmen, et al., 2025; Gast, Solla, & Kennedy, 2024; Mishra & Narayanan, 2016, 2021; Mittal & Narayanan, 2021; Saini & Narayanan, 2025; Tikidji-Hamburyan, Martinez, White, & Canavier, 2015; X. J. Wang & Buzsaki, 1996) Therefore, key questions that need to be addressed are as follows. How would the interplay between the heterogeneity in synaptic and intrinsic properties and sensory noise affect different aspects of the network performance? Could the network maintain robustness in grid-patterned firing across noise even when heterogeneities are introduced? Would the system exhibit stochastic resonance even when such heterogeneities are present?

In the context of heterogeneities in CAN models, neuronal resonance has been shown to serve as a stabilizing mechanism for heterogeneous CAN models (Mittal & Narayanan, 2021). Neuronal resonance is mediated by ion-channel conductances that introduce a slow negative feedback loop (Hutcheon & Yarom, 2000; Mishra & Narayanan, 2025; Mittal & Narayanan, 2021, 2024). There are several lines of evidence to support the occurrence of resonance in neurons in the medial entorhinal cortex (Canto & Witter, 2012; Erchova, Kreck, Heinemann, & Herz, 2004; L. M. Giocomo, Zilli, Fransen, & Hasselmo, 2007; Haas & White, 2002; Mittal & Narayanan, 2022; Pastoll, Ramsden, & Nolan, 2012). Future studies could therefore explore the possibility of whether intrinsic neuronal resonance might suppress the deleterious impact of noise through such a negative feedback loop. In addition, the slow time constants of these resonating conductances could also increase the time over which neurons could integrate self-motion inputs, which might also lead to an increase in accuracy in position estimate. Since navigation under realistic scenarios involves cluttered regions with several obstacles, object/landmark inputs from the lateral entorhinal cortex (Deshmukh & Knierim, 2011, 2013; Edvardsen, Bicanski, & Burgess, 2020) could be explored as an additional corrective influence.

Lastly, we incorporated sensory noise in our model and observed the occurrence of stochastic resonance in network performance. However, we incorporated additive white noise sources in our model, and there are studies that demonstrate that the color of noise plays a critical role as well (Gingl, Kiss, & Moss, 1995; Hänggi, Jung, Zerbe, & Moss, 1993; Nozaki, Mar, Grigg, & Collins, 1999; Nozaki & Yamamoto, 1998). Thus, future studies could look to examine the effect of changing the color of the noise or introducing multiplicative noise could alter the outcomes observed in our model.

## ACKNOWLEDGMENTS

The authors thank members of the cellular neurophysiology laboratory and Dr. Sarthak Chandra for helpful comments on a draft of this manuscript. This work was supported by the Wellcome Trust-DBT India Alliance (Senior fellowship to R. N.; IA/S/16/2/502727) and the Kishore Vaigyanik Protsahan Yojana (H. N.).

## Notes

### Competing Interest Statement

The authors have declared no competing interest.

### Summary of Updates

Several new additional results added. Analyses expanded.

## REFERENCES

Brandon, M. P., Bogaard, A. R., Libby, C. P., Connerney, M. A., Gupta, K., & Hasselmo, M. E. (2011). Reduction of theta rhythm dissociates grid cell spatial periodicity from directional tuning. Science, 332, 595–599.

Brunel, N., & Hakim, V. (1999). Fast global oscillations in networks of integrate-and-fire neurons with low firing rates. Neural Comput, 11, 1621–1671.

Burak, Y., & Fiete, I. R. (2009). Accurate path integration in continuous attractor network models of grid cells. PLoS Comput Biol, 5, e1000291.

Bush, D., & Burgess, N. (2014). A hybrid oscillatory interference/continuous attractor network model of grid cell firing. J Neurosci, 34, 5065–5079.

Cannon, S. C., Robinson, D. A., & Shamma, S. (1983). A proposed neural network for the integrator of the oculomotor system. Biol Cybern, 49, 127–136.

Canto, C. B., & Witter, M. P. (2012). Cellular properties of principal neurons in the rat entorhinal cortex. II. The medial entorhinal cortex. Hippocampus, 22, 1277–1299.

Cheung, A., Ball, D., Milford, M., Wyeth, G., & Wiles, J. (2012). Maintaining a cognitive map in darkness: the need to fuse boundary knowledge with path integration. PLoS Comput Biol, 8, e1002651.

Collins, J. J., Chow, C. C., & Imhoff, T. T. (1995a). Aperiodic stochastic resonance in excitable systems. Phys Rev E Stat Phys Plasmas Fluids Relat Interdiscip Topics, 52, R3321–R3324.

Collins, J. J., Chow, C. C., & Imhoff, T. T. (1995b). Stochastic resonance without tuning. Nature, 376, 236–238.

Dahmen, D., Hutt, A., Indiveri, G., Kennedy, A., Lefebvre, J., Mazzucato, L., Motter, A. E., Narayanan, R., Payvand, M., Planert, H., & Gast, R. (2025). How Heterogeneity Shapes Dynamics and Computation in the Brain. Neuron, In press, DOI: 10.1016/j.neuron.2025.1011.1023.

Deshmukh, S. S., & Knierim, J. J. (2011). Representation of non-spatial and spatial information in the lateral entorhinal cortex. Front Behav Neurosci, 5, 69.

Deshmukh, S. S., & Knierim, J. J. (2013). Influence of local objects on hippocampal representations: Landmark vectors and memory. Hippocampus, 23, 253–267.

Dong, L. L., & Fiete, I. R. (2024). Grid Cells in Cognition: Mechanisms and Function. Annual Review of Neuroscience, Vol 36, 47, 345–368.

Douglass, J. K., Wilkens, L., Pantazelou, E., & Moss, F. (1993). Noise enhancement of information transfer in crayfish mechanoreceptors by stochastic resonance. Nature, 365, 337–340.

Edvardsen, V., Bicanski, A., & Burgess, N. (2020). Navigating with grid and place cells in cluttered environments. Hippocampus, 30, 220–232.

Erchova, I., Kreck, G., Heinemann, U., & Herz, A. V. (2004). Dynamics of rat entorhinal cortex layer II and III cells: characteristics of membrane potential resonance at rest predict oscillation properties near threshold. The Journal of physiology, 560, 89–110.

Faisal, A. A., Selen, L. P., & Wolpert, D. M. (2008). Noise in the nervous system. Nat Rev Neurosci, 9, 292–303.

Fuhs, M. C., & Touretzky, D. S. (2006). A spin glass model of path integration in rat medial entorhinal cortex. J Neurosci, 26, 4266–4276.

Fyhn, M., Molden, S., Witter, M. P., Moser, E. I., & Moser, M. B. (2004). Spatial representation in the entorhinal cortex. Science, 305, 1258–1264.

Gardner, R. J., Hermansen, E., Pachitariu, M., Burak, Y., Baas, N. A., Dunn, B. A., Moser, M.-B., & Moser, E. I. (2022). Toroidal topology of population activity in grid cells. Nature, 1–6.

Gast, R., Solla, S. A., & Kennedy, A. (2024). Neural heterogeneity controls computations in spiking neural networks. Proc Natl Acad Sci U S A, 121, e2311885121.

Gingl, Z., Kiss, L., & Moss, F. (1995). Non-dynamical stochastic resonance: Theory and experiments with white and arbitrarily coloured noise. EPL (Europhysics Letters), 29, 191.

Giocomo, L. M. (2016). Environmental boundaries as a mechanism for correcting and anchoring spatial maps. The Journal of physiology, 594, 6501–6511.

Giocomo, L. M., Zilli, E. A., Fransen, E., & Hasselmo, M. E. (2007). Temporal frequency of subthreshold oscillations scales with entorhinal grid cell field spacing. Science, 315, 1719–1722.

Gluckman, B. J., Netoff, T. I., Neel, E. J., Ditto, W. L., Spano, M. L., & Schiff, S. J. (1996). Stochastic Resonance in a Neuronal Network from Mammalian Brain. Phys Rev Lett, 77, 4098–4101.

Haas, J. S., & White, J. A. (2002). Frequency selectivity of layer II stellate cells in the medial entorhinal cortex. Journal of neurophysiology, 88, 2422–2429.

Hafting, T., Fyhn, M., Molden, S., Moser, M. B., & Moser, E. I. (2005). Microstructure of a spatial map in the entorhinal cortex. Nature, 436, 801–806.

Hänggi, P., Jung, P., Zerbe, C., & Moss, F. (1993). Can colored noise improve stochastic resonance? Journal of Statistical Physics, 70, 25–47.

Hardcastle, K., Ganguli, S., & Giocomo, L. M. (2015). Environmental boundaries as an error correction mechanism for grid cells. Neuron, 86, 827–839.

Hasselmo, M. E., & Brandon, M. P. (2012). A model combining oscillations and attractor dynamics for generation of grid cell firing. Front Neural Circuits, 6, 30.

Hutcheon, B., & Yarom, Y. (2000). Resonance, oscillation and the intrinsic frequency preferences of neurons. Trends Neurosci, 23, 216–222.

Khona, M., & Fiete, I. R. (2022). Attractor and integrator networks in the brain. Nat Rev Neurosci, 23, 744–766.

Marder, E., & Taylor, A. L. (2011). Multiple models to capture the variability in biological neurons and networks. Nat Neurosci, 14, 133–138.

McDonnell, M. D., & Abbott, D. (2009). What is stochastic resonance? Definitions, misconceptions, debates, and its relevance to biology. PLoS Comput Biol, 5, e1000348.

Mishra, P., & Narayanan, R. (2016). Degenerate mechanisms mediate decorrelation and pattern separation in the dentate gyrus. In Society for Neuroscience Annual Meeting (Vol. Program No. 263.08). San Diego, USA.

Mishra, P., & Narayanan, R. (2021). Ion-channel regulation of response decorrelation in a heterogeneous multi-scale model of the dentate gyrus. Curr Res Neurobiol, 2, 100007.

Mishra, P., & Narayanan, R. (2025). The enigmatic HCN channels: A cellular neurophysiology perspective. Proteins, 93, 72–92.

Mittal, D., & Narayanan, R. (2021). Resonating neurons stabilize heterogeneous grid-cell networks. Elife, 10.

Mittal, D., & Narayanan, R. (2022). Heterogeneous stochastic bifurcations explain intrinsic oscillatory patterns in entorhinal cortical stellate cells. Proc Natl Acad Sci U S A, 119, e2202962119.

Mittal, D., & Narayanan, R. (2024). Network motifs in cellular neurophysiology. Trends Neurosci, 47, 506–521.

Moss, F. (1997). Stochastic resonance at the molecular level. Biophys J, 73, 2249–2250.

Navratilova, Z., Giocomo, L. M., Fellous, J. M., Hasselmo, M. E., & McNaughton, B. L. (2012). Phase precession and variable spatial scaling in a periodic attractor map model of medial entorhinal grid cells with realistic after-spike dynamics. Hippocampus, 22, 772–789.

Nozaki, D., Mar, D. J., Grigg, P., & Collins, J. J. (1999). Effects of colored noise on stochastic resonance in sensory neurons. Physical Review Letters, 82, 2402.

Nozaki, D., & Yamamoto, Y. (1998). Enhancement of stochastic resonance in a FitzHugh-Nagumo neuronal model driven by colored noise. Physics Letters A, 243, 281–287.

Pastoll, H., Ramsden, H. L., & Nolan, M. F. (2012). Intrinsic electrophysiological properties of entorhinal cortex stellate cells and their contribution to grid cell firing fields. Frontiers in neural circuits, 6, 17.

Pollock, E., Desai, N., Wei, X.-x., & Balasubramanian, V. (2018). Dynamic self-organized error-correction of grid cells by border cells. arXiv preprint arXiv:1808.01503.

Rathour, R. K., & Narayanan, R. (2019). Degeneracy in hippocampal physiology and plasticity. Hippocampus, 29, 980–1022.

Ribrault, C., Sekimoto, K., & Triller, A. (2011). From the stochasticity of molecular processes to the variability of synaptic transmission. Nat Rev Neurosci, 12, 375–387.

Russell, D. F., Wilkens, L. A., & Moss, F. (1999). Use of behavioural stochastic resonance by paddle fish for feeding. Nature, 402, 291–294.

Saini, S., & Narayanan, R. (2025). Degeneracy Explains Diversity in Interneuronal Regulation of Pattern Separation in Heterogeneous Dentate Gyrus Networks. Function (Oxf), 6.

Samsonovich, A., & McNaughton, B. L. (1997). Path integration and cognitive mapping in a continuous attractor neural network model. J Neurosci, 17, 5900–5920.

Savelli, F., & Knierim, J. J. (2019). Origin and role of path integration in the cognitive representations of the hippocampus: computational insights into open questions. J Exp Biol, 222.

Savelli, F., Yoganarasimha, D., & Knierim, J. J. (2008). Influence of boundary removal on the spatial representations of the medial entorhinal cortex. Hippocampus, 18, 1270–1282.

Simonotto, E., Riani, M., Seife, C., Roberts, M., Twitty, J., & Moss, F. (1997). Visual Perception of Stochastic Resonance. Physical Review Letters, 78, 1186–1189.

Solstad, T., Boccara, C. N., Kropff, E., Moser, M. B., & Moser, E. I. (2008). Representation of geometric borders in the entorhinal cortex. Science, 322, 1865–1868.

Tikidji-Hamburyan, R. A., Martinez, J. J., White, J. A., & Canavier, C. C. (2015). Resonant Interneurons Can Increase Robustness of Gamma Oscillations. J Neurosci, 35, 15682–15695.

Wang, R., & Kang, L. (2022). Multiple bumps can enhance robustness to noise in continuous attractor networks. PLoS Comput Biol, 18, e1010547.

Wang, X. J., & Buzsaki, G. (1996). Gamma oscillation by synaptic inhibition in a hippocampal interneuronal network model. J Neurosci, 16, 6402–6413.

