## Supplementary Figures S1-S4 for "Stochastic resonance in the impact of afferent sensory noise on grid-patterned firing and path integration in a continuous attractor network"

### Supplementary material

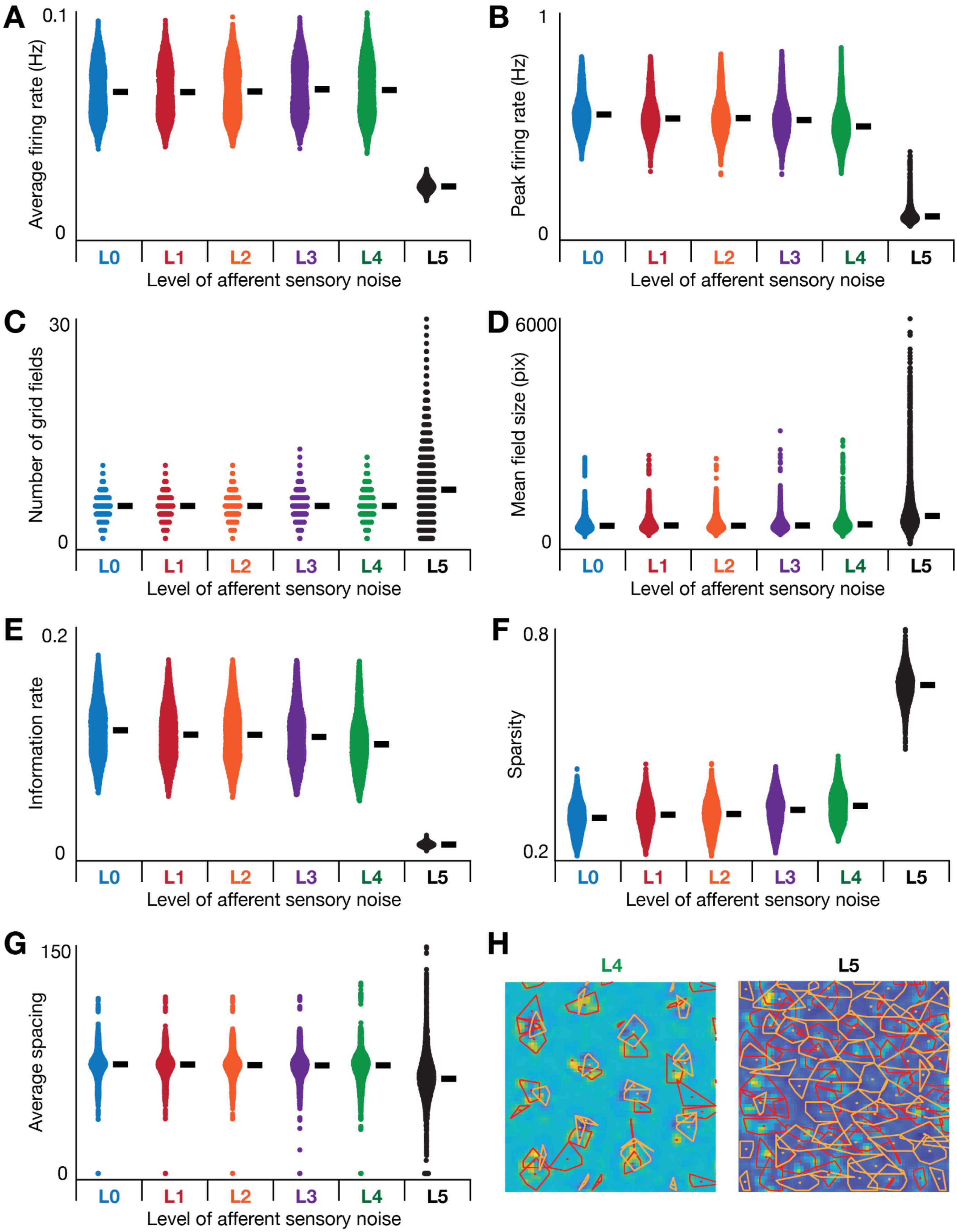

**Supplementary Figure S1. Measurements associated with grid-patterned firing with different levels of afferent noise.** (*A–G*) Bee-swarm plots of different properties of the 3600 neurons in the network, obtained with all 6 levels of afferent noise. *(H)* Example activity difference maps with the convex hull for maxima (red) and minima (blue) at the L4 and L5 levels of noise. The contours depict the inability to identify convex hulls for centroid computation towards obtaining network pattern flow velocity.

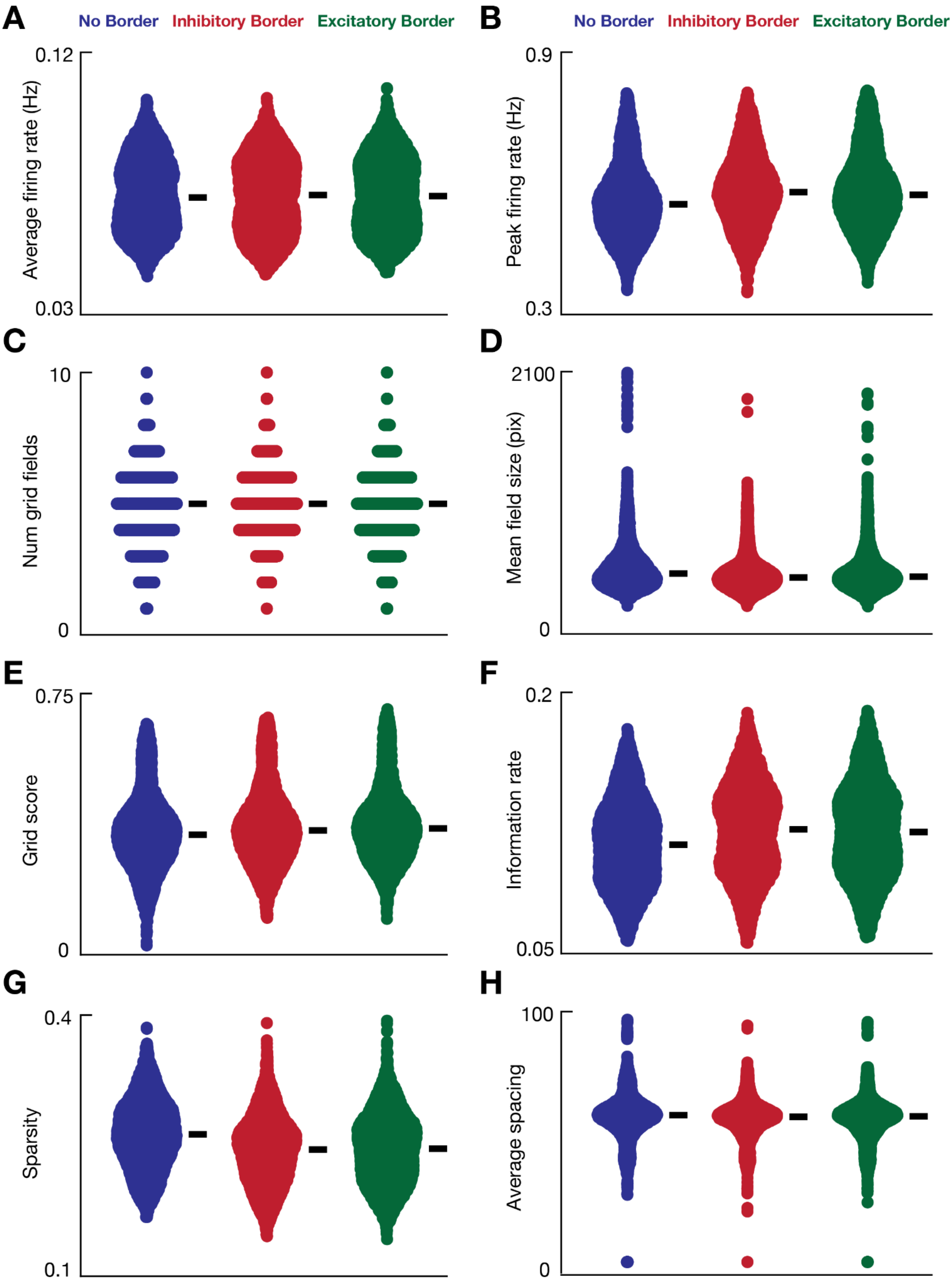

**Supplementary Figure S2. Introduction of border cells introduced modest changes to measurements related to grid-patterned firing.** Bee-swarm plots of the different properties of the 3600 neurons in the network generated for a representative trajectory, with colors representing the type of border input to the network. The black bars represent the respective medians.

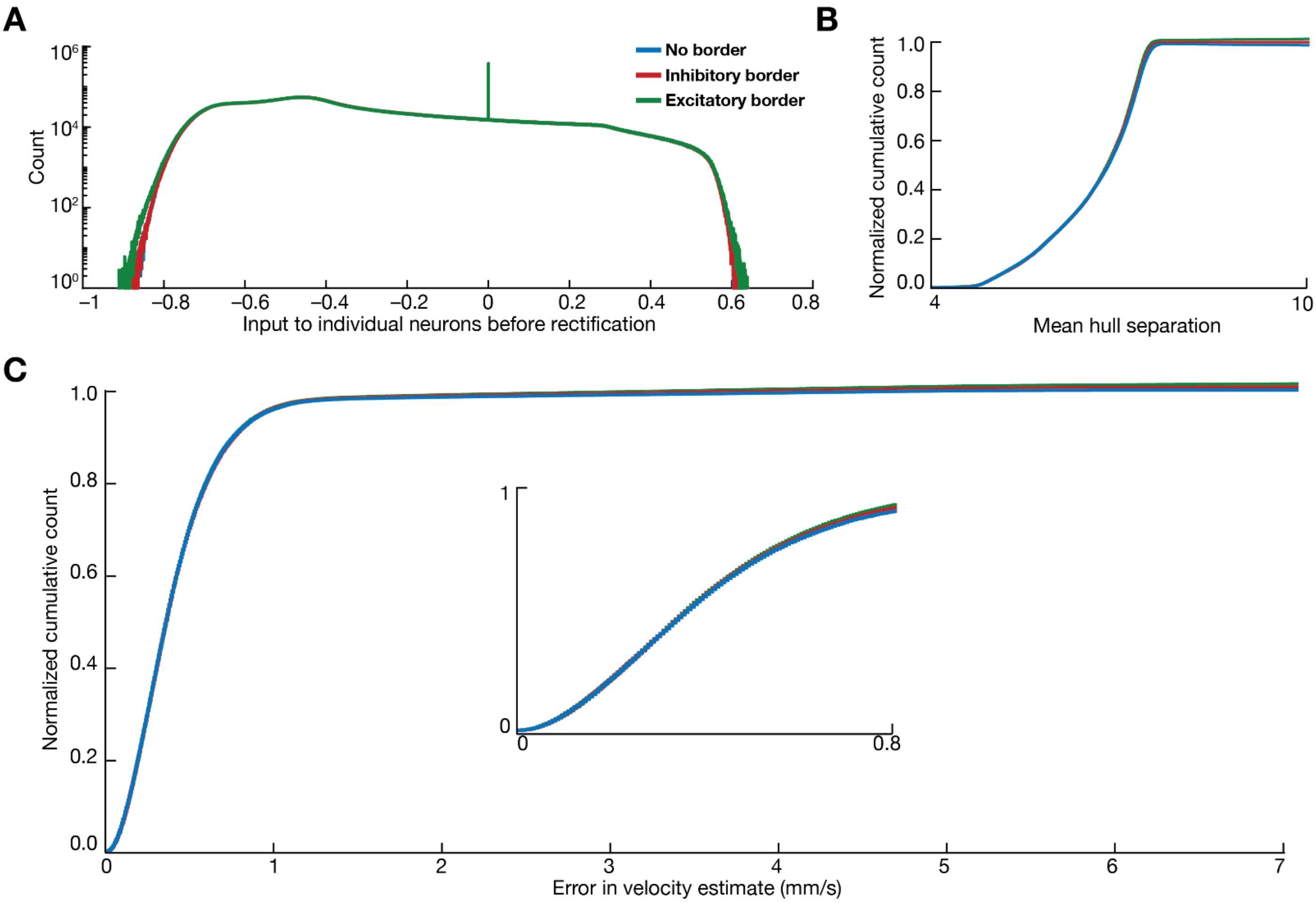

**Supplementary Figure S3. Introducing border cells yielded modest impact on input distribution, hull separation, and velocity estimates.** *(A)* Distribution of pre-rectification inputs for the 3600 neurons of the network for a representative run for the 3 different types of border inputs introduced to the network. *(B)* The distribution of the mean hull separation when the 3 different types of borders are introduced. *(C)* The distribution of the error in velocity estimate when the 3 different types of borders are introduced. *Inset*, A zoomed-in version of the same distribution spanning low error values.

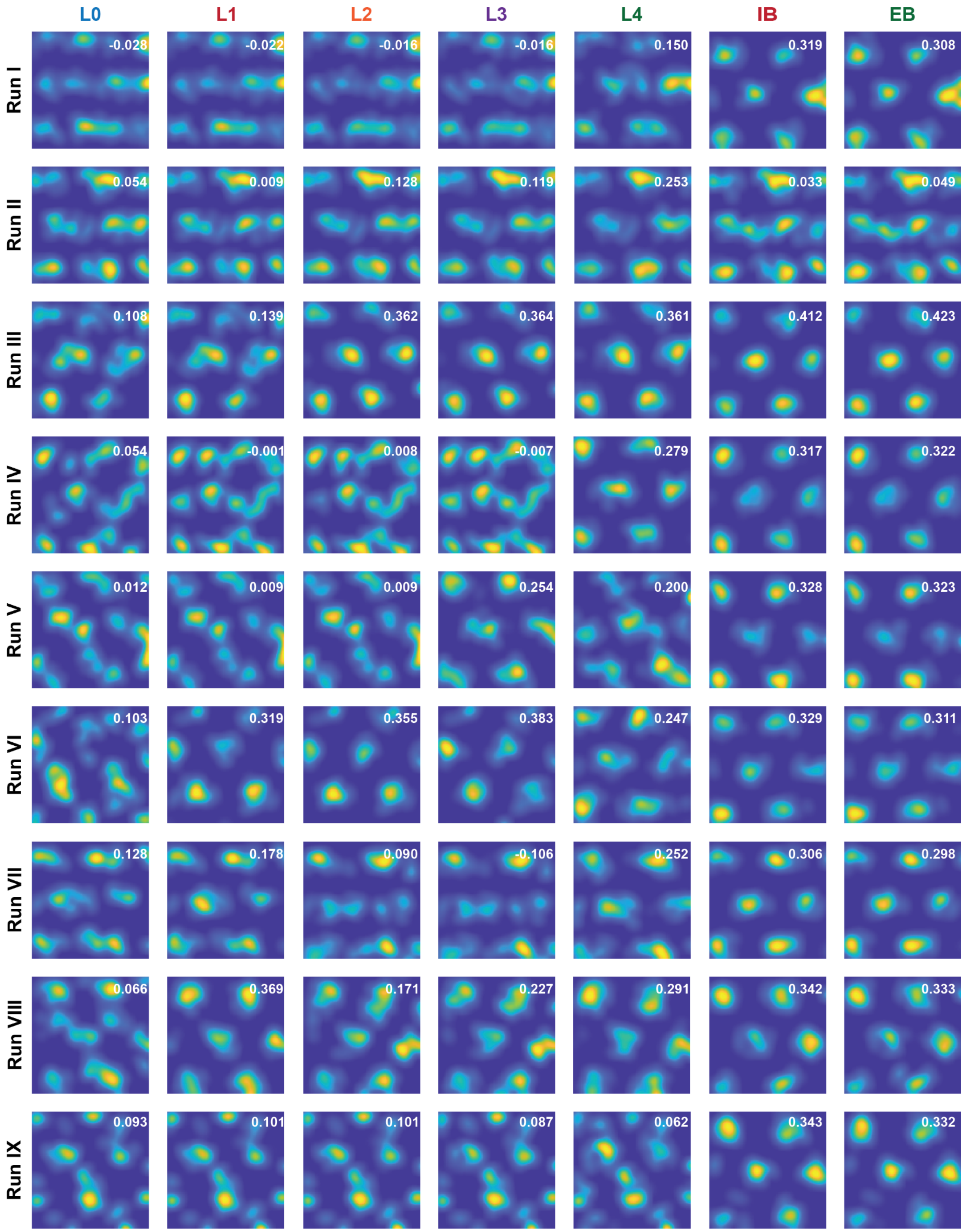

**Supplementary Figure S4. Sensory noise and border inputs are beneficial for the generation of grid fields.** Example rate maps of a single cell for different noise levels and border cell inputs are shown. Each row represents a single trajectory while the values in white represent the mean grid score.

**Supplementary Table S1: Results of statistical Wilcoxon signed rank test for data shown in Figure 1*B*.**

| | L0 | L1 | L2 | L3 | L4 |
|---|---|---|---|---|---|
| **L1** | $2.18\times10^{-28}$ | | | | |
| **L2** | $2.87\times10^{-70}$ | $4.22\times10^{-55}$ | | | |
| **L3** | $1.46\times10^{-06}$ | $9.45\times10^{-36}$ | $1.25\times10^{-118}$ | | |
| **L4** | $4.62\times10^{-107}$ | $1.27\times10^{-153}$ | $5.34\times10^{-200}$ | $8.49\times10^{-162}$ | |
| **L5** | 0.00 | 0.00 | 0.00 | 0.00 | 0.00 |
